# Fatty acid dysregulation in the anterior cingulate cortex of depressed suicides with a history of child abuse

**DOI:** 10.1101/2021.06.21.449337

**Authors:** Kelly Perlman, Raphaël Chouinard-Watkins, Arnaud Tanti, Giulia Cisbani, Massimiliano Orri, Gustavo Turecki, Richard P Bazinet, Naguib Mechawar

**Author notes:** Corresponding Author: Naguib Mechawar, Douglas Mental Health University Institute, 6875 Boulevard LaSalle, Montréal, QC, H4H 1R3.

## Abstract

Child abuse (CA) strongly increases the lifetime risk of suffering from major depression and predicts an unfavorable course for the illness. Severe CA has been associated with a specific dysregulation of oligodendrocyte function and thinner myelin sheaths in the human anterior cingulate cortex (ACC) white matter. Given that myelin is extremely lipid-rich, it is plausible that these findings may be accompanied by a disruption of the lipid profile that composes the myelin sheath. This is important to explore since the composition of fatty acids (FA) in myelin phospholipids can influence its stability, permeability, and compactness. Therefore, the objective of this study was to quantify and compare FA concentrations in postmortem ACC white matter in the choline glycerophospholipid pool (ChoGpl), a key myelin phospholipid pool, between adult depressed suicides with a history of CA (DS-CA) matched depressed suicides without CA (DS) and healthy non-psychiatric controls (CTRL). Total lipids were extracted according to the Folch method and separated into respective classes using thin-layer chromatography. FA methyl esters from the ChoGpl fraction were quantified using gas chromatography. Our analysis revealed a strong age-related decrease in most FAs, and specific effects of CA in FAs from the arachidonic acid synthesis pathway, which was further validated with RNA-sequencing data. Furthermore, the concentration of most FAs was found to decrease with age. By extending the previous molecular level findings linking CA with altered myelination in the ACC, these results provide further insights regarding white matter alterations associated with early-life adversity.

## Introduction

Child abuse (CA) is a major public health problem. In 2016, it was estimated that, worldwide, one billion children between 2 and 17 years of age experienced maltreatment in the previous year^1^. Overall, victims of CA present an increased burden of disease, both physical and psychological^2^. In particular, CA strongly increases the lifetime risk of suffering from major depressive disorder (MDD) and predicts an unfavorable course for the illness as well as poorer response to treatment^3^. Furthermore, CA comprises 54% of the population attributable risk (PAR) for depression and 67% of the PAR for suicide attempts^4^. Importantly, CA is the primary preventable risk factor in the development of mental illness^4^. It has been posited that the experience of CA modifies neural development such that the brain becomes more susceptible to psychopathology of MDD and suicide^5^, making it such a potent risk factor. Converging evidence in both animal and human studies suggest that the process of myelination contributes to this CA-induced vulnerability^5, 6^.

Numerous CA-related myelin findings have been identified in the anterior cingulate cortex (ACC) a key limbic brain region involved in emotional regulation, processing of social pain, error detection, attention, cognitive control, reward-based decision making and learning^7, 8^. Our group has previously reported that in the ACC, there is an impairment in the myelin program of both the epigenome and the transcriptome, and that the myelin sheath around small diameter axons is significantly thinner in postmortem brain samples from depressed suicides with a history of CA compared to matched samples from depressed suicides and non-psychiatric controls with no history of CA^9^.

The myelin sheath is highly enriched in lipids, with proportions estimated at 70% to 85%, in comparison to the typical membrane which is comprised of about 50% lipids^10^. Of the lipid pool, approximately 40% is constituted of phospholipids, of which nearly one third is represented by the choline glycerophospholipid pool (ChoGpl)^11, 12^. It is estimated that of the ChoGpl pool, 90% is comprised of phosphatidylcholine (PC), with the remaining fraction being choline plasmalogens^13^. In phosphatidylcholine, a choline head group is esterified to the phosphate group at the Sn-3 position, and two fatty acids, usually one saturated (at the Sn-1 position) and one unsaturated (at the Sn-2 position), are attached to a glycerol backbone through ester linkages^14^. PC is a key structural and functional phospholipid in myelin and is a precursor for a number of cell signalling molecules^12^.

Lipid studies of MDD have revealed that lipid dysregulation may be part of the pathophysiology of the illness and that it may be a promising target for treatment, often with a focus on plasma omega-3 FAs^15–17^. Several pre-clinical and postmortem human brain studies of MDD versus control individuals have identified differences in FA between groups, mostly when considering total lipids, though results are highly heterogeneous^18–23^. These inconclusive results may in part be explained by various factors in the experimental design of these studies. Notably, most studies in the field have examined total FAs, which can obfuscate any lipid class specific FA findings. Moreover, there has often been a lack of information regarding the grey and white matter composition of the tissue samples, which is consequential because there are documented differences in FA composition between grey and white matter^24^. Lastly, most studies have reported brain FA content as molar percent rather than absolute concentration, and the lack of an internal standard makes it impossible to control for the size of the FA pools.

The objective of this study was to quantify and compare ChoGpl FA concentrations in postmortem ACC white matter, between depressed suicides with a history of severe CA (DS- CA) compared to matched depressed suicides without CA (DS) and healthy controls (CTRL). Our analysis revealed significant differences in concentrations between groups in the FAs that compose the arachidonic acid synthesis pathway. Furthermore, the concentration of most FAs was found to be inversely correlated with age.

## Methods

### Samples

This study was approved by the Douglas Mental Health University Institute’s Research Ethics Board. Brain samples were provided by the Douglas-Bell Canada Brain Bank (DBCBB; http://douglasbrainbank.ca). All brains are donated to the Suicide section of the DBCBB by familial consent through the Quebec Coroner’s Office. Suicide and sudden-death control brains undergo a validated process known as psychological autopsy to retrieve phenotypic information^25^. Histories of abuse were measured using adapted Childhood Experience of Care and Abuse scale interviews - cases with maximum severity levels for abuse having occurred before the age of 15 were included, as in previous work^9^.

As displayed in Table 1, frozen dorsal ACC white matter was obtained from a total of 101 individuals (15-85 years old) divided into three groups: male and female depressed suicides with (DS-CA; n=39) or without (DS; n=36) a history of severe CA and matched sudden-death controls with no history of psychiatric nor neurological conditions (CTRL; n=26). Samples were dissected from coronal brain sections of the left hemisphere by expert brain bank staff, as described previously^9^.

**Table 1.** Subject information.

|  | **CTRL** | **DS-CA** | **DS** |
| --- | --- | --- | --- |
| **N total** | 26 | 39 | 36 |
| **N female** | 7 | 11 | 5 |
| **Age (yrs)\*** | 46.46 ± 19.86 | 40.49 ± 14.38 | 50.19 ± 11.34 |
| **pH** | 6.48 ± 0.28 | 6.50 ± 0.34 | 6.48 ± 0.35 |
| **PMI (h)** | 42.30 ± 27.70 | 53.24 ± 27.62 | 57.04 ± 28.43 |

Age, pH, and PMI data are expressed as mean ± standard deviation. To test for significant differences between groups for age and pH, a one-way ANOVA was used with a cut-off of $p < 0.05$ . The data for PMI failed the Shapiro-Wilks test for normality, so a Kruskal-Wallis test was used with a cut off of $p < 0.05$ . Age: $p = 0.02$ , pH: $p = 0.94$ , PMI: $p = 0.098$ .

### Lipid extraction and FA quantification

A minimum of 30 mg of tissue was used per sample. Total lipids were extracted according to the Folch method (2:1 ratio of chloroform to methanol), homogenized in a glass douncer^26^. Lipids were separated into respective classes using thin-layer chromatography (TLC) on silica H-plates. The mobile phase was 30:9:25:6:18 chloroform:methanol:2-propanol:0.25% KCl: triethylamine (v/v/v/v/v). The plate was run until the solvent front reached 2 cm from the top. Once dried, the plate was sprayed with with 0.1% (w/v) 8-anilino-1-naphthalene sulfonic acid and bands were viewed under ultraviolet light. The ChoGpl fraction was scraped off of the TLC plate and transmethylated using 14% BF3 in methanol with incubation for 1 hour at 100°C. FA methyl esters from ChoGpl were quantified using a Varian 430 gas chromatograph (Bruker, Billerica, MA, USA). C17:0 was used as an internal standard and concentrations (measured in μg/g) of each FA were derived from the area under the curve (AUC) of its respective peak on the chromatogram. Relative percentage was calculated based on the individual and total FA concentrations. A graphical summary of this methodology is illustrated in Figure 1.

**Figure 1.**
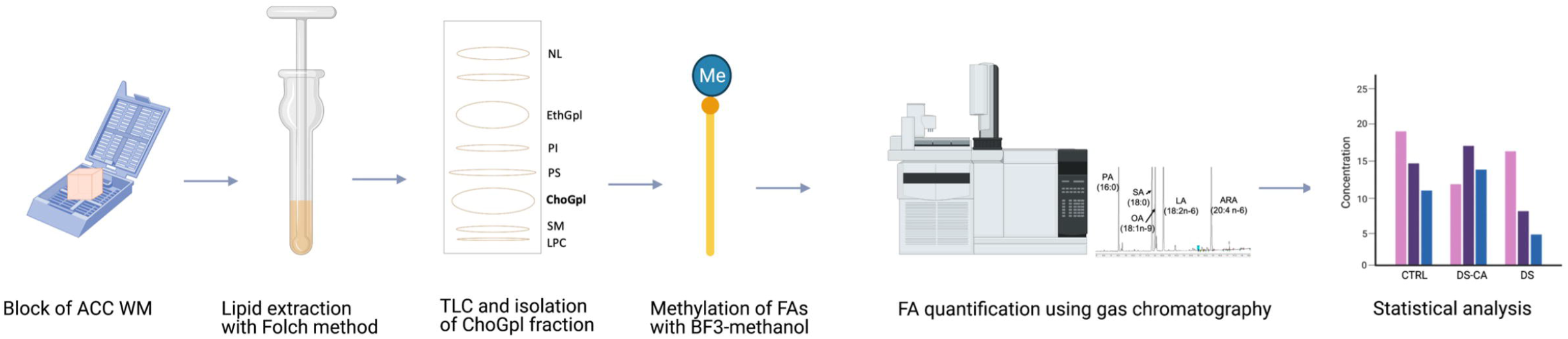
Graphical overview of FA quantification methodology. NL: neutral lipids (e.g., cholesterol), EthGpl: ethanolamine glycerophospholipids, PI: phosphatidylinositol, PS: phosphatidylserine, ChoGpl: choline glycerophospholipids, SM: sphingomyelin, LPC: lysophosphatidylcholine. Figure was created using biorender.com.

### Gene expression

The RNA sequencing data was previously generated in the Lutz, Tanti et al. (2017) study, which examined the transcriptome of ACC grey matter in DS-CA versus CTRL groups. Briefly, RNA was exacted from frozen tissue, and only samples with an RNA integrity number (RIN) greater than 5 was used. Libraries were built using TrueSeq Standed Total RNA Sample Preparation and Ribo-Zero gold kits. All libraries were sequenced at the Genome Quebec Innovation Center on the Illumina HiSEquation 2000 platform, and each library with a sequencing depth of approximately 62 million reads. Differential expression was calculated using the DESeq2 package^27^.

### Statistics

All statistics were computed in R. We modelled FA concentration and relative percentage as a function of group with age, postmortem interval (PMI), pH, and sex (encoded using a dummy variable) as covariates. Standard errors were computed using nonparametric bootstrap, (k=10000) with the ANOVA.boot function of the lmboot package. Using this procedure, the coefficients of interest of the models were estimated 10000 times in datasets using resampling, which generates an empirical distribution of coefficients from which robust standard errors can be estimated. This procedure achieve more robust and accurate estimations than parametric statistics in samples of limited size^28^. P-values were corrected for multiple comparisons using the Benjamini-Hochberg (BH) method, which yielded q-values (q < 0.05 was considered significant). Group-by-age interactions were tested but none reached statistical significance after BH correction and as such were not included in the models. Estimated marginal means for groups were calculated across all covariates and plotted with the emmeans package. Pearson correlation was used to assess the relationship between FA concentrations and were considered significant at p<0.05.

## Results

### Sample exclusions

Concentration and relative percentage data were generated with samples from 101 subjects (Table 1). One DS sample was excluded due to technical artifacts. As can be seen on the principal component analysis plot for concentration (Supplementary Figure 1), this sample is separated from all others across PC1 (i.e., the axis corresponding to the linear transformation that accounts for the most variance in the data), which clearly makes it an extreme outlier. We were not able to rerun this sample due to tissue unavailability. Furthermore, one CTRL sample was excluded from the between-group analysis because medical reports obtained later noted that this subject had been prescribed antidepressants, though no psychiatric diagnosis was noted.

### Between group modelling

We compared 20 FAs between groups, as well as a total FA quantification for the concentration metric. FAs with concentrations significantly different between groups after BH correction were: C18:2n-6, C20:3n-6, C20:4n-6, (2m, 2o, 2p). Our analyses were not sufficiently powered to detect significant differences between individual pairs of groups in post-hoc tests. However, the estimated marginal means (EMM) for groups displayed highest concentration in the DS-CA group for all significant FAs. There did appear to be a trend, in which the EMMs several of the FAs (including the total FA metric) were higher in the DS-CA group, though this is solely descriptive (Figure 2). For each model with a significant group term, we calculated the percentage of variance explained by each significant factor (Supplementary Table 2).

**Figure 2.**
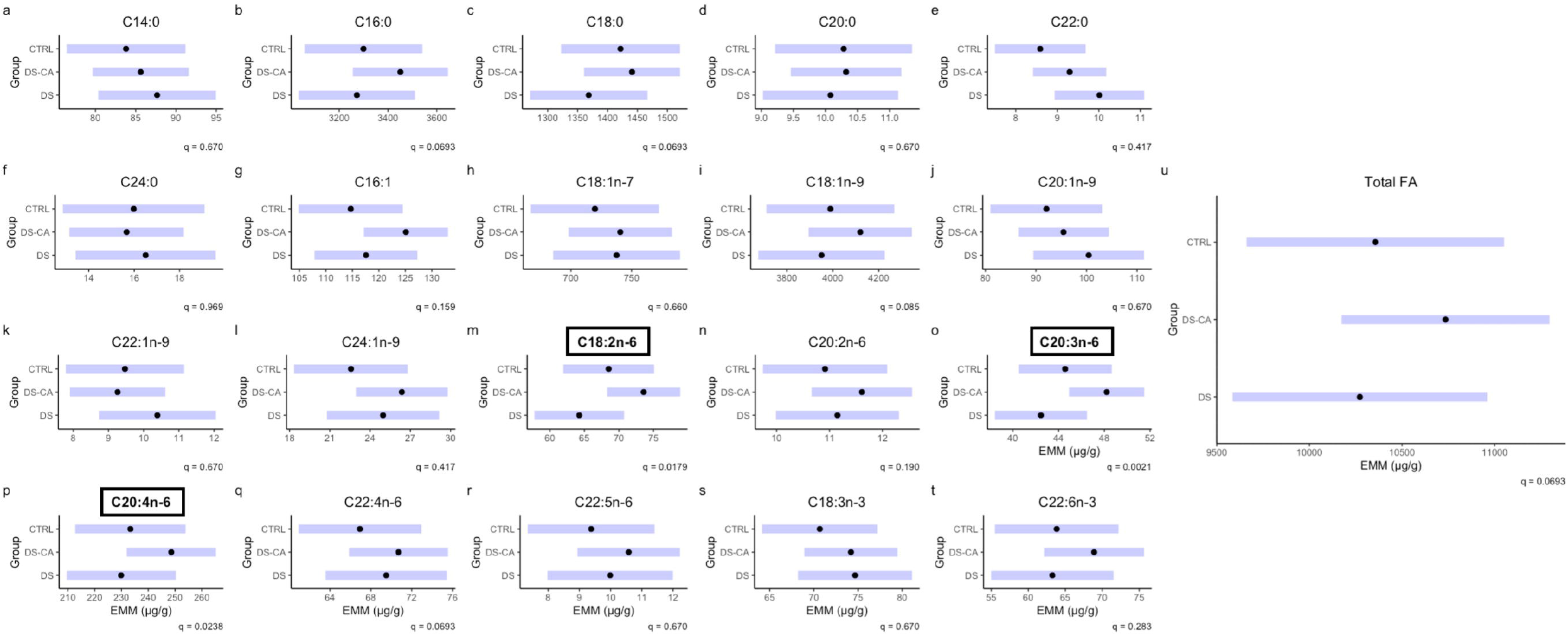
Concentration of fatty acids in choline glycerophospholipids between groups. Black dots represent the estimated marginal means for group and purple bars represent the 95% confidence intervals based on linear models for concentration of FA as a function of group with age, PMI, pH, and sex as covariates. Subplots with boxed titles in bold font indicate significance between groups with q < 0.05. Concentrations are represented in μg/g.

When considering the group differences between FAs in relative percentage, we found that no FA reached significance after correction with BH (Figure 3), despite highly similar patterns for C18:2n-6, C20:3n-6, C20:4n-6, as in concentration. For reference, Supplementary Table 1 lists the relative percentage of individual FAs in ascending order.

**Figure 3.**
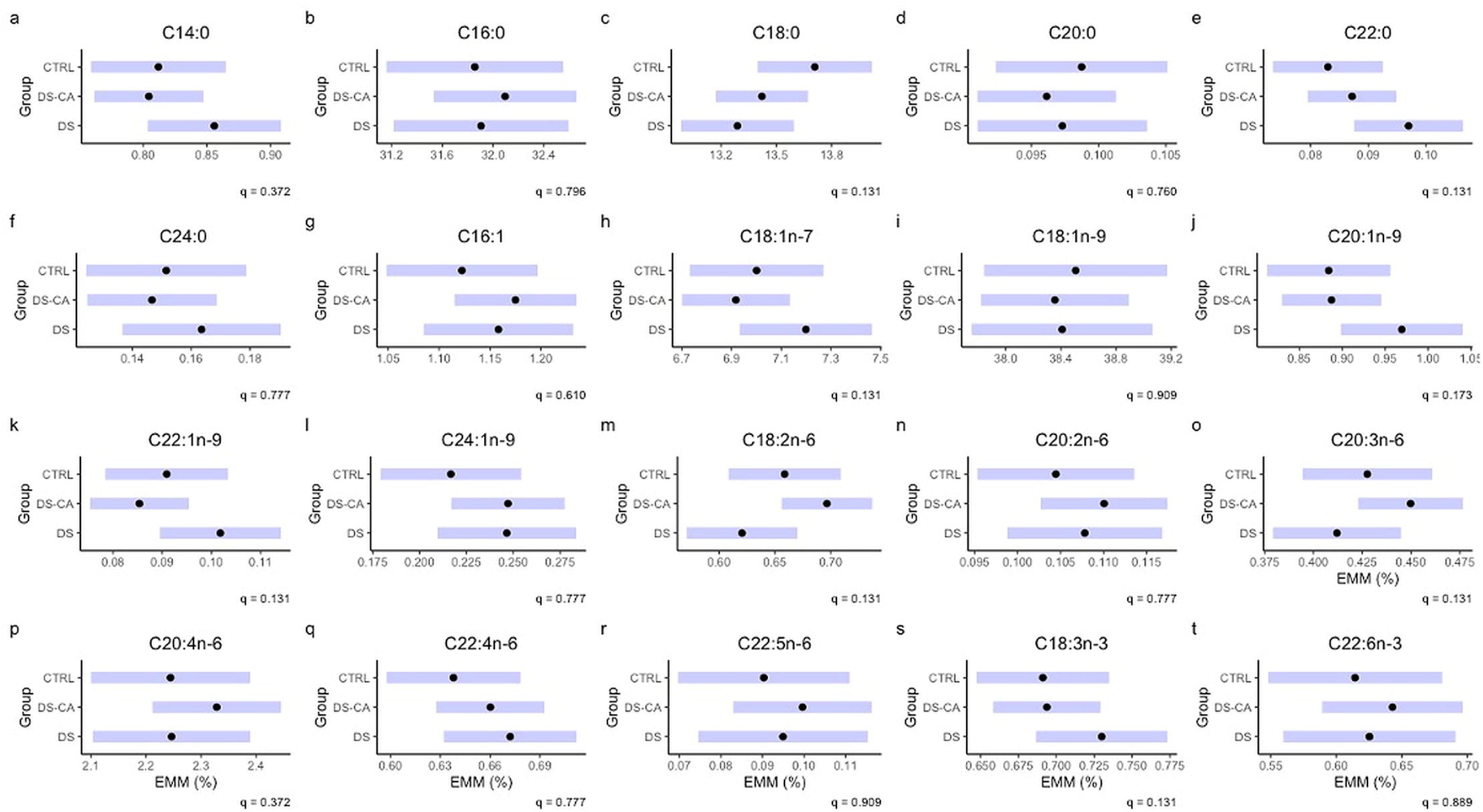
Relative percentage of fatty acids in choline glycerophospholipids between groups. Black dots represent the estimated marginal means for group and the purple bars represent the 95% confidence intervals based on linear models for concentration of FA as a function of group with Age, PMI, pH, and sex as covariates. The group term did not reach significance in any relative percentage model after correction for multiple comparisons.

### Age correlations

Upon examining the relationships between FA quantities and the covariates of interest, we observed that age was the covariate most strongly correlated (in terms of magnitude of the correlation coefficient) compared to the others, in particular when considering concentration (Supplementary Figure 2A). Given the wide age span of the subjects included in this study, we were able to study the association between age and FA levels from adolescence through old age. Since group differences were not being considered here, we added back the CTRL subject excluded due to medication history.

Pearson correlations demonstrated significant relationships for 14 out of 20 FAs (plus the total FA metric) such that age was inversely correlated with concentration of these FAs (Figure 4a). The FAs with the highest correlation coefficient were C20:3n-6 (r = −0.44), 20:4n-6 (r = −0.48) and C22:4n-6, (r = −0.57). Only these FAs with the strongest negative correlations for concentration appeared to be reflected as decreased across the age span for relative percentage (Figure 4b). Furthermore, none of the 4 FAs that were significantly directly correlated with age in terms of relative percentage (C16:1, C18:1n-7, C18:2n-6, C18:3n-3) reached significance when correlating age and concentration (Figure 4b). As such, it is likely that these FAs may contribute a greater proportion of the total FA pool later in life as their concentrations remain fairly constant while others are decreasing.

**Figure 4.**
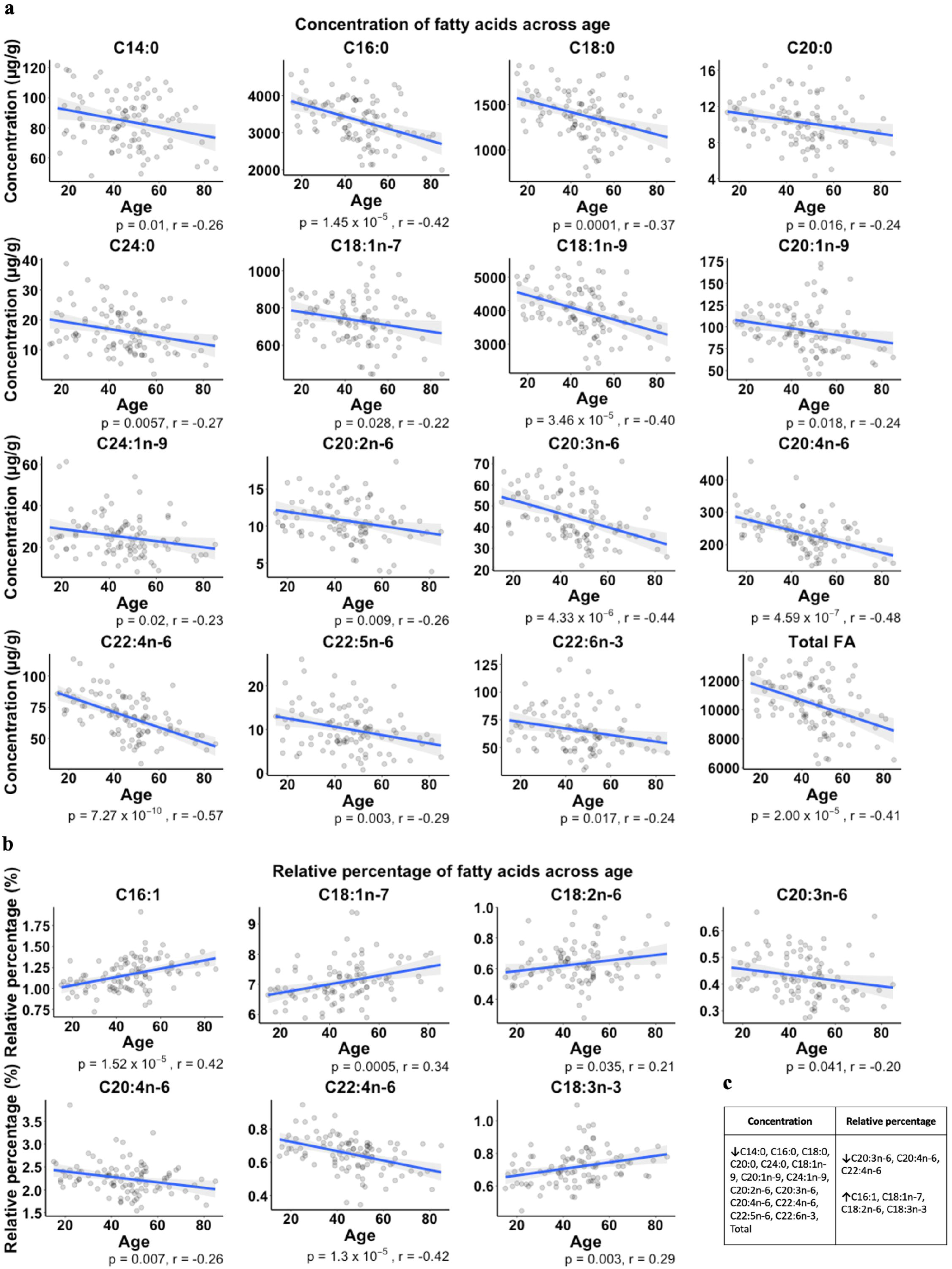
Significant correlations between age and fatty acid quantitative metrics. Significant Pearson correlations between a) age (years) and concentrations (μg/g) of FAs and b) age (yrs) and relative percentage. P-values and r-values are listed under each subplot. A summary table of these significant correlations is presented in c) in which negative correlation is represented by a downward arrow and positive correlation is represented by an upward arrow.

### Transcriptomics

Given that the FAs that showed significant differences between groups were all involved in the arachidonic acid (AA) synthesis pathway, we aimed to assess whether this finding was reflected in the respective gene expression profiles. We obtained a list of genes in the AA metabolic process (GO:0019369) and looked for the intersection of those genes with the genes from the ACC RNA-seq with nominal p-value < 0.05. Table 2 lists the gene names and symbols, along with the corresponding nominal p-values, fold changes and log fold changes. We observed that key genes involved in AA metabolism, including PLA2G4A and PTGS2 (also known as COX-2), were significantly differentially expressed between DS-CA and CTRL groups in the ACC grey matter.

**Table 2.**
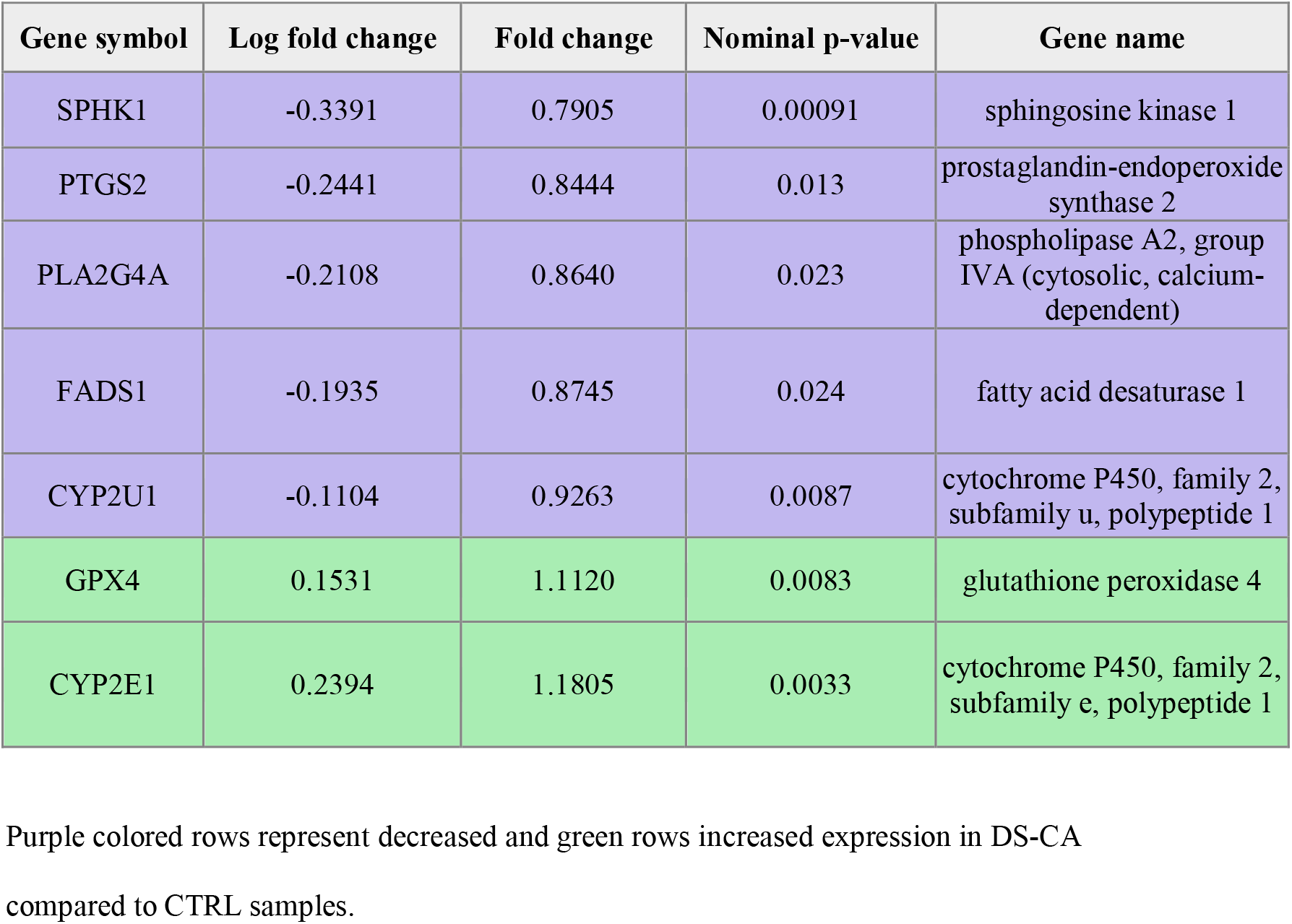
Genes involved in arachidonic acid metabolism with significant p-values.

## Discussion

In this study of the quantification of ChoGpl FAs in human ACC, our key findings are that 1) the concentration of key FAs are dysregulated between DS-CA, DS, and CTRL groups; 2) the concentration of most FAs as well as the total FA concentration tends to decrease with age; and 3) concentration appears to be a more robust metric as compared to relative percentage for this type of research.

This FA dysregulation observed between groups may be related to our previous finding that the myelin sheath of small caliber axons in the ACC white matter are thinner in the DS-CA group^9^. The composition of FAs in myelin phospholipids has been demonstrated to influence its stability^10^, permeability^29,30^ and compactness^31^. Therefore, one potential mechanism for the altered myelin sheath properties observed in CA^9, 32^ could be a disruption of the lipid profile that composes the membrane, resulting in changes to properties such as membrane fluidity, which can lead to alterations in structural integrity with consequences on efficient information flow. This is speculated because FAs with higher concentrations in the DS-CA group (at least descriptively) comprise the majority of ChoGpl according to abundance (Figure 2, Supplementary Table 1) and accordingly reflected in the total concentration (Figure 2u), possibly pointing to a greater quantity of overall ChoGpl in this group. There is also transcriptomic evidence to support this view, since genes coding for key enzymes in PC synthesis such as choline kinase alpha and Choline/ethanolaminephosphotransferase 1 are differentially expressed between the DS-CA and CTRL groups (Supplementary Table 3). The mechanisms linking these FA changes to larger scale morphology changes require further study, with consideration of the other lipid classes in myelin.

To our knowledge, this is the first postmortem study to examine the potential impact of CA on brain FAs. In rats, it was demonstrated that a maternal separation paradigm induced a shift in the plasma PUFAs towards a more pro-inflammatory profile, with an increased concentration of AA in the maternally separated group^33^. In humans, previous work has investigated the relationship between plasma PUFA levels in young adults, early life trauma, and depressive symptoms, which revealed that low omega-3 FA levels and a history of trauma are both associated with persistence of depressive symptoms, but not correlated to each other^34^. However, plasma and brain FA levels do not appear to be correlated^35^, emphasizing the importance of human postmortem brain tissue with well-documented histories of maltreatment for obtaining a comprehensive picture of lipid-related changes associated with early-life adversity.

The AA synthesis-related FA and gene expression findings suggest a possible dysregulation of inflammatory processes in the DS-CA group. While the directionality of these findings appears contradictory, it must be highlighted that the RNA-seq data were generated from grey matter while the FA results from white matter. It remains unclear, however, whether these genes are differentially regulated between cortical compartments. A more likely explanation of the apparent discrepancy between gene expression and FA levels may be that elevated levels of all the quantified FAs in the AA synthesis pathway may have driven compensatory changes in gene expression. Also, while AA is a precursor to inflammatory mediators, it is implicated in other biological processes including ion channel regulation, cell signalling, neurotransmitter release, and even neurogenesis^36, 37^, so the precise functional role it may play in this context requires further investigation. Furthermore, since dysregulation of the HPA axis is a consistent finding in individuals with a history of childhood maltreatment^38^, it may be worth investigating the relationship between glucocorticoids as an inhibitor of phospholipase A2 (PLA2)^39^ and any adaptive responses thereof.

Our findings concerning age are consistent with several previous postmortem studies demonstrating decreasing FA levels as a function of age^40–42^. In a study of total FAs from cerebral cortex homogenates, Carver and colleagues found decreasing concentrations of FAs, in particular omega-6 FAs, across ages 18 to 88 years^41^. Hancock et al. found decreasing levels of AA and adrenic acid (C22:4n-6) in combined phospholipids across ages 18 to 104 years in both mitochondrial and microsomal membrane fractions of hippocampus. Notably, AA and adrenic acid were the FAs with the largest negative correlation (according to r value) in our data. Moreover, administration of PUFAs has been shown to ameliorate cognitive deficits and even mild cognitive impairment that occurs in old age^43^, which may reflect the beneficial effects of compensating for the observed loss over time.

This study also serves to illustrate that the concentration metric can be useful to prevent overlooking or mischaracterizing findings that may result from considering only relative percentage. For example, relative percentage cannot detect global upregulation or downregulation owing to the fact that it cannot control for the size of the FA pools. This observation has also been made in a study of mild cognitive impairment and Alzheimer’s disease, such that no changes were observed when using a relative percentage metric but were identified using concentration data.^35^ Absolute metrics obtained through the use of an internal standard should become common practice in the field of FA quantification in brain tissue.

This study is not without limitations. Firstly, we considered the ChoGpl pool as a whole, so we were unable to obtain a ratio of PC to choline plasmalogens for each subject. Despite the vast majority of the pool being comprised of PC, it is possible that the abundance ratio differed across subjects. This may be an especially important distinction given that plasmalogens have been documented to protect myelin against damage from reactive oxygen species^44^, and oxidative stress is thought to be a mechanism linking early life stress to the development of later psychopathology^45^. Notably, we examined only the ChoGpl pool of phospholipids, and it was not possible to get a complete picture of any global dysregulation without studying other lipid classes. Furthermore, due to the low number of females in our sample, we were likely unable to reliably detect potential sex-specific differences. In addition, we extracted ChoGpl from blocks of ACC white matter, and therefore do not know if our findings are driven by the FAs of a specific subcellular compartment (e.g., the mitochondrial membrane). Another limitation comes from the fact that the available RNA-seq data only compared DS-CA and CTRL samples, so the differentially expressed genes could not be specifically associated with CA without further validation with samples from all 3 groups. However, this transcriptomic data was used purely as a resource to explore potential links between FA differences and gene expression in relevant biological pathways. Finally, since we do not know the source of these FAs, we cannot comment on whether the group differences in FA concentration are reflective of alterations in synthesis, transport, and/or dietary intake. A future direction of this research is to probe dietary intake by assessing lipogenesis using carbon 13 isotopes in gas chromatography-combustion-isotope ratio mass spectrometry^46^.

In summary, we have established that the concentrations of various ChoGpl FAs are altered between DS-CA, DS, and CTRL groups, and that they tend to decline across the lifespan. These findings warrant further investigation, in particular to establish the underlying mechanisms and to determine how they may contribute to the neurobiological vulnerability to psychopathology resulting from early-life adversity.

## Acknowledgements

This work was supported by a CIHR Project Grant to NM and RPB. KP holds an FRQS scholarship and both GC and RCW hold CIHR postdoctoral fellowships. RPB hold the Canada Research Chair in Brain Lipid Metabolism. The Douglas-Bell Canada Brain Bank (DBCBB) is partly funded by a Healthy Brains for Healthy Lives (CFREF) Platform Grant to GT and NM. The DBCBB is also funded by the Réseau Québécois sur le suicide, le troubles de l’humeur et les troubles associés (FRQS).

