## Supplementary Material for "Fatty acid dysregulation in the anterior cingulate cortex of depressed suicides with a history of child abuse"

Supplementary Figure 1 – page 2

Supplementary Figure 2 – page 3

Supplementary Table 1 – page 4

Supplementary Table 2 – page 5

Supplementary Table 3 – page 6

**Supplementary Figure 1**
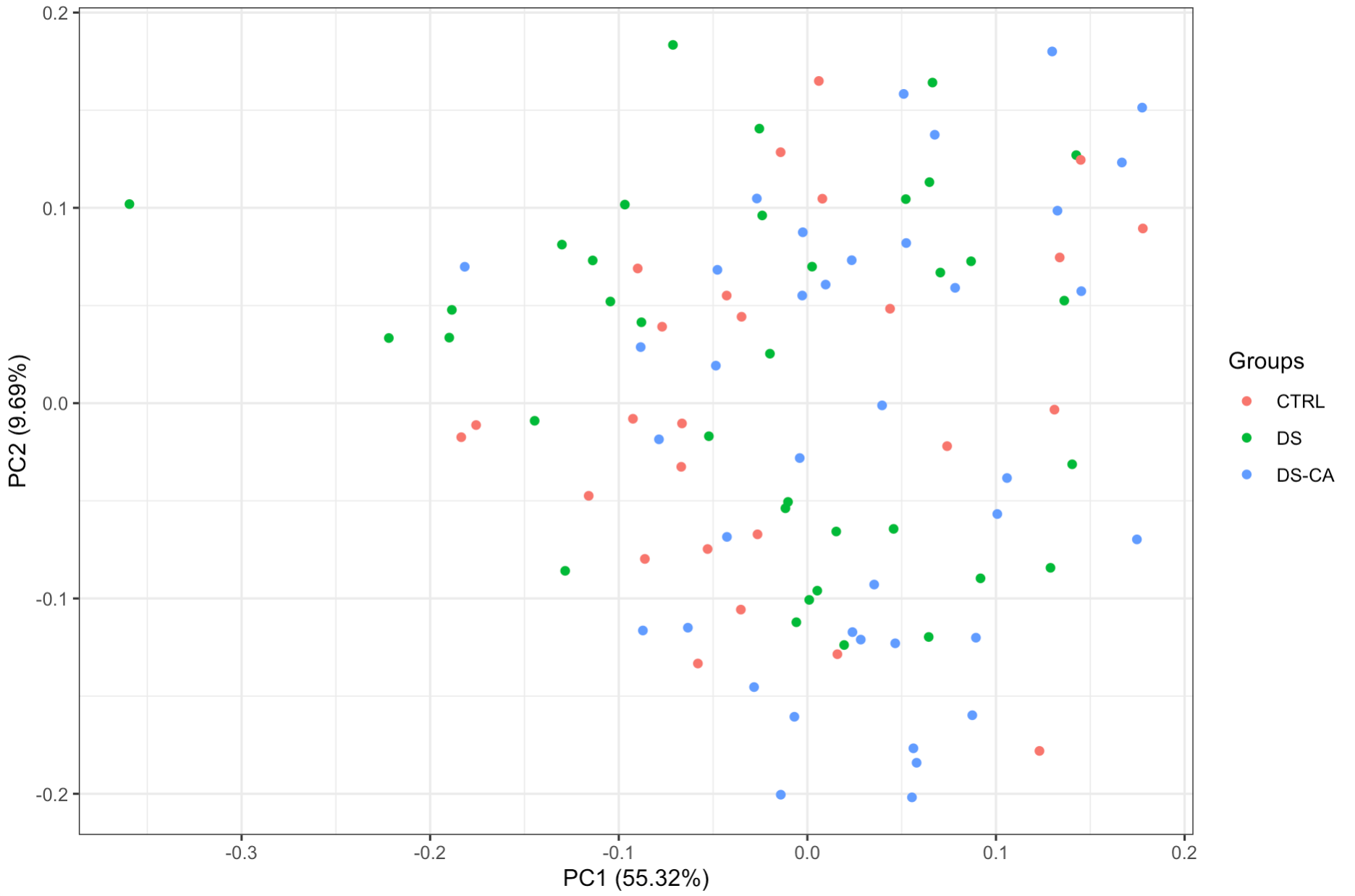

Supplementary Figure 1 – Scatter plot of the first two principal components from a PCA of the FA concentration data, with each point representing one subject. Each point is colored by subject group and the percentage displayed next to each axis label represents the variance explained by each component. A clear outlier from the DS group can be seen along PC1, indicated by an arrow.

**Supplementary Figure 2**

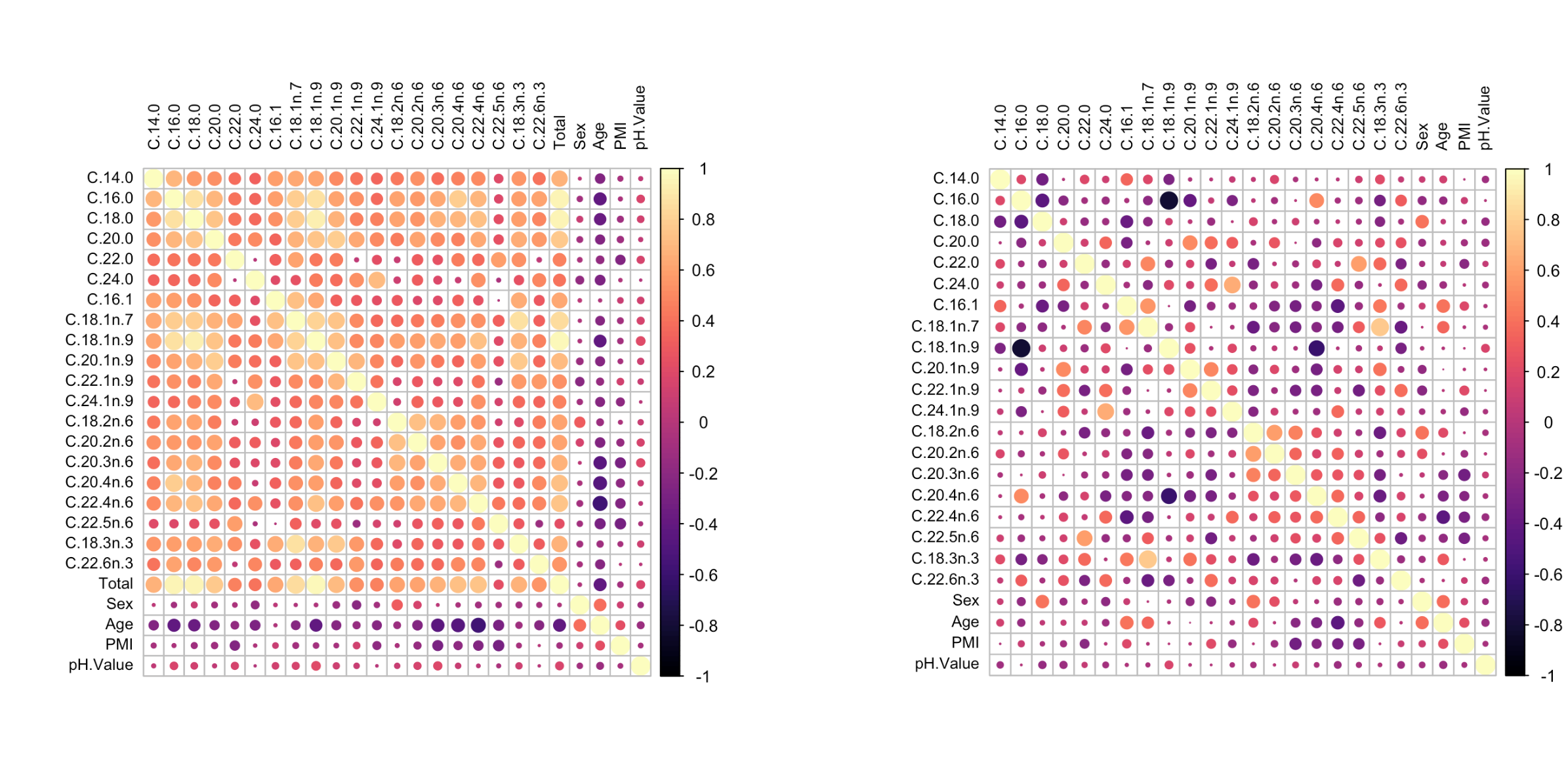

**a**

**b**

**Concentration**

**Relative percentage**

Supplementary Figure 2- Correlation plots for sex, age, PMI, pH, and all FAs for both a) concentration and b) relative percentage metrics. The size and color of each circle is representative of the correlation coefficient.

**Supplementary Table 1** – Relative percentages by FA across all subjects in ascending order

| FA | Mean (%) | SD (%) |
| --- | --- | --- |
| C22:0 | 0.0899 | 0.0230 |
| C22:1n-9 | 0.0955 | 0.0309 |
| C22:5n-6 | 0.0980 | 0.0489 |
| C20:0 | 0.0986 | 0.0146 |
| C20:2n-6 | 0.1027 | 0.0214 |
| C24:0 | 0.1568 | 0.0620 |
| C24:1n-9 | 0.2413 | 0.0845 |
| C20:3n-6 | 0.4287 | 0.0818 |
| C18:2n-6 | 0.6285 | 0.1270 |
| C22:6n-3 | 0.6310 | 0.1486 |
| C22:4n-6 | 0.6510 | 0.1043 |
| C18:3n-3 | 0.7160 | 0.1060 |
| C14:0 | 0.8166 | 0.1215 |
| C20:1n-9 | 0.9221 | 0.1664 |
| C16:1 | 1.1648 | 0.1843 |
| C20:4n-6 | 2.2496 | 0.3459 |
| C18:1n-7 | 7.0812 | 0.6499 |
| C18:0 | 13.2791 | 0.7968 |
| C16:0 | 32.1272 | 1.5739 |
| C18:1n-9 | 38.4217 | 1.4874 |

Mean and standard deviation of the relative percentages for all subjects (excluding the DS outlier) are displayed in ascending order with respect to the mean.

**Supplementary Table 2**- Percentage of variance explained by factors in the significant FA models

| **FA** | **Group (variance explained)** | **Covariate (variance explained)** |
| --- | --- | --- |
| C18:2n-6 | 11.48% | Sex: 6.73% |
| C20:3n-6 | 13.92% | Age: 12.18% |
| C20:4n-6 | 9.28% | Age: 16.90% |

Linear models of the format lm(FA ~ group + covariate) were created for the FAs with significant p-values for group in the bootstrapped models. Any significant covariates for those FAs were included in the linear model, and the percentage the overall variance explained by each factor was calculated from that model.

**Supplementary Table 3 –** Significantly differentially expressed genes involved in phosphatidylcholine metabolism

| **Gene symbol** | **Log fold change** | **Fold change** | **Nominal p-value** | **Gene name** |
| --- | --- | --- | --- | --- |
| PLA2G4A | -0.2981914 | 0.8132713 | 0.00353957 | Cytosolic phospholipase A2 |
| LCAT | -0.271347 | 0.8285456 | 0.00097269 | Phosphatidylcholine-sterol acyltransferase |
| LPCAT2 | -0.2565997 | 0.83705846 | 0.00107114 | Lysophosphatidylcholine acyltransferase 2 |
| PLA2G5 | -0.2173768 | 0.86012793 | 0.00428426 | Calcium-dependent phospholipase A2 |
| MBOAT1 | -0.1823435 | 0.88127033 | 0.00100037 | Lysophospholipid acyltransferase 1 |
| ENPP2 | -0.1822277 | 0.88134102 | 0.0031028 | Ectonucleotide pyrophosphatase/phosphodiesterase family member 2 |
| CHKA | -0.14409 | 0.90495003 | 0.00218141 | Choline kinase alpha |
| PLB1 | -0.09873 | 0.93385469 | 0.00373475 | Phospholipase B1 |
| FABP3 | 0.22571474 | 1.16935643 | 0.00206479 | Fatty acid-binding protein 3 |
| ABHD3 | 0.26856898 | 1.20461237 | 1.8017E-05 | Phospholipase ABHD3 |
| MBOAT2 | 0.27677509 | 1.21148378 | 2.3491E-05 | Lysophospholipid acyltransferase 2 |
| CEPT1 | 0.31916572 | 1.24760887 | 2.7806E-05 | Choline/ethanolaminephosphotransferase 1 |

List of genes in the phosphatidylcholine metabolic process (GO: 0046470) and that intersect with those from the ACC RNA-seq with nominal p-value < 0.05. Purple colored rows represent decreased and green rows increased expression in DS-CA compared to CTRL samples.
